# The role of microRNAs in understanding sex-based differences in Alzheimer’s disease

**DOI:** 10.1101/2023.08.24.554586

**Authors:** Jaime Llera-Oyola, Héctor Carceller, Zoraida Andreu, Marta R. Hidalgo, Irene Soler-Sáez, Fernando Gordillo, Antonio Porlan, Macarena Pozo-Morales, Beatriz Roson, Maria de la Iglesia-Vayá, Roberta Mancuso, Franca R. Guerini, Akiko Mizokami, Francisco García-García

## Abstract

**Background:** The incidence of Alzheimer’s disease (AD) - the most frequent cause of dementia - is expected to increase as life expectancies rise across the globe. While sex-based differences in AD have previously been described, there remain uncertainties regarding any association between sex and disease-associated molecular mechanisms. Studying sex-specific expression profiles of regulatory factors such as microRNAs (miRNAs) could contribute to more accurate disease diagnosis and treatment.

**Methods:** A systematic review identified five studies of microRNA expression in AD patients that incorporated information regarding the biological sex of samples in the Gene Expression Omnibus repository. A differential microRNA expression analysis was performed, considering disease status and patient sex. Subsequently, results were integrated within a meta-analysis methodology, with a functional enrichment of meta-analysis results establishing an association between altered miRNA expression and relevant Gene Ontology terms.

**Results:** Meta-analyses of miRNA expression profiles in blood samples revealed the alteration of sixteen miRNAs in female and twenty-two miRNAs in male AD patients. We discovered nine miRNAs commonly overexpressed in both sexes, suggesting a shared miRNA dysregulation profile. Functional enrichment results based on miRNA profiles revealed sex-based differences in biological processes; most affected processes related to ubiquitination, regulation of different kinase activities, and apoptotic processes in males, but RNA splicing and translation in females. Meta-analyses of miRNA expression profiles in brain samples revealed the alteration of six miRNAs in female and four miRNAs in male AD patients. We observed a single underexpressed miRNA in female and male AD patients (*hsa-miR-767-5p*); however, the functional enrichment analysis for brain samples did not reveal any specifically affected biological process.

****Conclusions**:** Sex-specific meta-analyses supported the detection of differentially expressed miRNAs in female and male AD patients, highlighting the relevance of sex-based information in biomedical data. Further studies on miRNA regulation in AD patients should meet the criteria for comparability and standardization of information.

**Highlights:**

- Deregulation of miRNA expression profiles occurs in a tissue- and sex-specific manner in AD patients
- Meta-analysis of blood samples revealed a partial overlapping pattern of altered miRNA expression in female and male AD patients
- Functional enrichment based on AD-associated miRNA expression profiles in blood samples reveals sex-based differences: RNA splicing and translation in female AD patients and ubiquitination, regulation of different kinase activities, and apoptotic process in male AD patients
- Links between AD development and miRNA expression in brain tissue also demonstrate the influence of sex

**Plain English Summary:** Alzheimer’s disease (AD) - a neurodegenerative disease mainly affecting older patients - is characterized by cognitive deterioration, memory loss, and progressive incapacitation in daily activities. While AD affects almost twice as many females as males, and cognitive deterioration and brain atrophy develop more rapidly in females, the biological causes of these differences remain poorly understood. MicroRNAs (miRNAs) regulate gene expression and impact a wide variety of biological processes; therefore, studying the differential expression of miRNAs in female and male AD patients could contribute to a better understanding of the disease. We reviewed studies of miRNA expression in female and male AD patients and integrated results using a meta-analysis methodology and then identified those genes regulated by the altered miRNAs to establish an association with biological processes. We found sixteen (females) and twenty-two (males) miRNAs altered in the blood of AD patients. Functional enrichment revealed sex-based differences in the affected altered biological processes - protein modification and degradation and cell death in male AD patients and RNA processing in female AD patients. A similar analysis in the brains of AD patients revealed six (females) and four (males) miRNAs with altered expression; however, our analysis failed to highlight any specifically altered biological processes. Overall, we highlight the sex-based differential expression of miRNAs (and biological processes affected) in the blood and brain of AD patients.

## Background

Alzheimer’s disease (AD) is a common progressive neurodegenerative disease that causes dementia in the older population; however, ∼5%-10% of all AD cases start to develop in people under 65 (early onset AD) [1]. AD incidence is estimated to triple by 2050 [2–4], representing a global challenge of increasing impact for public health systems. Cognitive deterioration and the loss of memory and social skills characterize AD symptomatology, which culminates in total dependency on others and death [5]. AD risk factors include age and a family history of AD; the latter mainly involves early-onset AD and the presence of mutations in the amyloid precursor protein (*APP*), presenilin-1 (*PSEN1*), or presenilin-2 (*PSEN2*) genes [6]. Genome-wide association studies have also identified risk-inducing mutations in late-onset AD, with affected genes involved in pathways that include cholesterol/lipid metabolism, inflammation/the immune system, and endosome cycling [7]. Pathological hallmarks of AD include the presence of amyloid plaques composed of aggregated β-amyloid peptide (Aβ), which form as a consequence of Aβ overproduction/insufficient removal, and neurofibrillary tangles composed of hyperphosphorylated tau, which extend throughout brain regions during disease progression [8–10]. Aβ accumulation leads to excitotoxicity, inflammation, and oxidative stress [8] due to microglial cell overactivation, which contributes to synaptic impairment [8–10], while hyperphosphorylated tau affects cytoskeletal stability, altering the trafficking of postsynaptic receptors and axonal transport. In addition, multiple neurotransmission systems (such as the acetylcholine, serotonin, and glutamate systems) display impairments due to inflammatory events; these effects produce alterations in memory, neuroplasticity, and excitotoxicity [9,11].

Research into potential sex-based differences in AD has described an increased prevalence and incidence in females [12], with increased life expectancy and socio-economic factors as a partial explanation [13], while numerous studies have also described the more rapid cognitive decline and atrophic rate in females [14–17]. However, sex-based differences in molecular profiles that may explain these differences and sex-specific AD biomarkers remain undescribed. Characterizing the sex-based expression profiles of microRNAs (miRNAs) could provide insight into the disease-associated deregulation of multiple biological processes. Current diagnostic techniques for AD, including the analysis of cerebrospinal fluid-resident biomarkers and neuroimaging analysis, remain of limited clinical potential due to their invasive nature or high cost [18,19]; therefore, the identification of novel biomarkers such as miRNAs may allow for the development of more affordable, non-invasive, and highly-sensitive techniques. The quantification of circulating miRNAs in the blood represents a promising non-invasive tool that could facilitate diagnosis and tailored interventions in AD patients for various reasons: i) miRNAs are small non-coding RNA molecules that regulate the expression of genes at post-transcriptional levels; ii) miRNA expression is conserved, temporal, and tissue-specific; iii) miRNAs influence the onset and pathology of AD as they play a role in Aβ metabolism, tau function and immunoinflammatory responses; iv) sex-based differential expression of miRNAs has been previously described in AD and other neurological diseases; and v) miRNAs may be regulated by sex hormones and display a higher density on the X chromosome [20][21][20][22][20][23,24][25–27]. These data suggest that evaluating sex-specific miRNA patterns will improve clinical outcomes in AD patients.

We conducted a systematic review and meta-analysis of miRNA expression studies in the blood and brain of female and male AD patients. A range of previous studies had integrated data from numerous sources to characterize sex-based differences in the human transcriptome [28–33]; however, to our knowledge, this study represents the first meta-analysis of microRNA in AD patients to provide a better understanding of the sex-related molecular mechanisms underlying the disease. We found consensus AD-associated miRNAs in both tissues and sex-specific miRNA expression signatures (especially in blood samples), potentially unveiling novel sex-based biomarkers for AD. Finally, we functionally characterized the effects of miRNAs with altered expression profiles in blood samples, describing sex-specific biological processes that become altered in AD patients.

## Methods

### Study selection via systematic review

A systematic review following the Preferred Reporting Items for Systematic Reviews and Meta-Analyses (PRISMA) guidelines [34] was performed, searching for the studies of microRNA expression in human AD patients (2010-2022) on the Gene Expression Omnibus (GEO) and ArrayExpress databases [35,36]. The keyword selected for the search was “Alzheimer’s disease” and the results were filtered by the following inclusion criteria: i) dataset type: “non-coding RNA profiling by array” or “non-coding RNA profiling by high throughput sequencing”, and ii) organism: “Homo sapiens”.

Exclusion criteria applied to the identified studies were: i) studies not related to AD, ii) studies without the sex information of each patient, iii) experimental designs different from AD patients versus controls, iv) transcriptomic studies not focused on miRNAs, v) studies on organisms other than humans, and vi) studies for which expression data was not accessible.

### Bioinformatic workflow

Our approach to expression data analysis comprised a pipeline for every study selected: i) data acquisition, ii) normalization and preprocessing, iii) exploratory analysis, and iv) differential expression analysis. The differential expression results were then integrated into a meta-analysis for each sex and tissue and the functional enrichment analysis of the meta-analysis results. All analyses were performed using the R language 4.1.2 [37], and the packages required to carry out the analyses were deposited in the Zenodo repository (http://doi.org/10.5281/zenodo.8385733).

### Data acquisition, normalization, and preprocessing

The normalized expression matrix and the sample information of each selected study were downloaded from the GEO database. Standard nomenclature for sex and health condition labels of each sample were reviewed and used to facilitate further analyses and comparisons. Only control or AD samples were selected in studies containing additional experimental groups.

For miRNA nomenclature, all features were annotated with miRBase v22 IDs [38], and mature miRNAs were filtered. The highest expression value was preserved for repeated miRNAs. Then, the minimum expression value was added to matrices to eliminate negative values, and log2 transformation was applied to those expression values not previously transformed.

### Exploratory analysis of individual studies

Data were explored using several graphical representations to provide an overview and identify potential anomalies in the data of each study: the proportion of patients by condition and sex and the expression data distribution with boxplots. Furthermore, potential categorical aggregations of samples associated with the experimental conditions were assessed via hierarchical clustering and principal component analysis (PCA).

### Differential expression analysis

Analyzing the differential expression of miRNAs used a linear model to assess the effects of AD on female and male patients using two comparisons: i) (AD female - Control female) and ii) (AD male - Control male). This analysis was conducted with the limma package for R [28]. A transformation from discrete to continuous data was made for RNA-sequencing studies using the “voom” function of the limma package, thus allowing the linear model construction. According to the linear model, differences in expression levels could be determined under the studied conditions for each miRNA analyzed and each comparison. The Benjamini & Hochberg (BH) method [39] was applied to adjust p-values for multiple comparisons.

### Meta-analysis

The differential expression analysis was integrated for individual studies with a meta-analysis approach, grouping the studies by tissue (brain and blood). For each group of studies, the meta-analysis was applied to the results of previously proposed comparisons (female and male), resulting in four meta-analyses (brain female, blood female, brain male, and blood male).

A random-effects meta-analysis methodology was selected (the DerSimonian and Laird approach [40]), which considers the expected heterogeneity of the studies involved. Meta-analyses were conducted using the metafor R package following a series of data processing steps [41]. Those miRNAs not present in at least two integrated studies were removed from the meta-analysis. For the remaining miRNAs, the logarithm of the fold change (LogFC) and its standard error computed in each study were combined to calculate the observed expression across all studies. The confidence interval for each LogFC calculated was adjusted for multiple comparisons with the BH method, and those miRNAs with adjusted p-value < 0.05 were considered significantly affected by AD.

### Functional enrichment

A functional enrichment methodology was carried out on the transcriptomic profile of each of the four meta-analyses to establish an association between miRNAs and their potentially functional effects. It was necessary to annotate the miRNAs with the genes on which they exert their regulation function to connect those affected genes with terms linked to them in the Gene Ontology (GO) [42]. The multiMiR package [43] allowed the identification of the genes targeted by each miRNA analyzed. The methodology described by García-García [44] and using functions present in the mdgsa package [45] were used to elaborate a ranked list of genes targeted by miRNAs, which were subsequently associated with their linked GO terms. The associations between genes and GO terms were downloaded from the Biomart database [46]. The statistical significance values of the altered functions were adjusted using the BH method, and those with an adjusted p-value < 0.05 were considered significantly affected.

### Metafun-AD-miRNA web tool

All data and results generated in the various steps of the meta-analyses are freely available on the Metafun-AD-miRNA platform (http://bioinfo.cipf.es/metafun-AD-miRNA) to any user, allowing the confirmation of obtained results and the exploration of other results of interest. This easy-to-use resource is divided into different sections: i) the summary of analysis results in each phase, followed by detailed results for the ii) exploratory analysis, iii) meta-analysis, and iv) functional profiling for each meta-analysis. The user can interact with the web tool through graphics and tables and explore information associated with specific miRNAs, genes, or biological functions.

## Results

We have organized the results of this study into four sections: i) the selection of studies from the systematic review, ii) the individual exploratory analysis carried out on each study, iii) the differential miRNA expression profiles from several comparisons, and iv) integration of differential miRNA expression results with a meta-analysis approach and the functional enrichment developed within a Gene Set Enrichment Analysis (GSEA) methodology (**Figure 1**).

**Figure 1.**
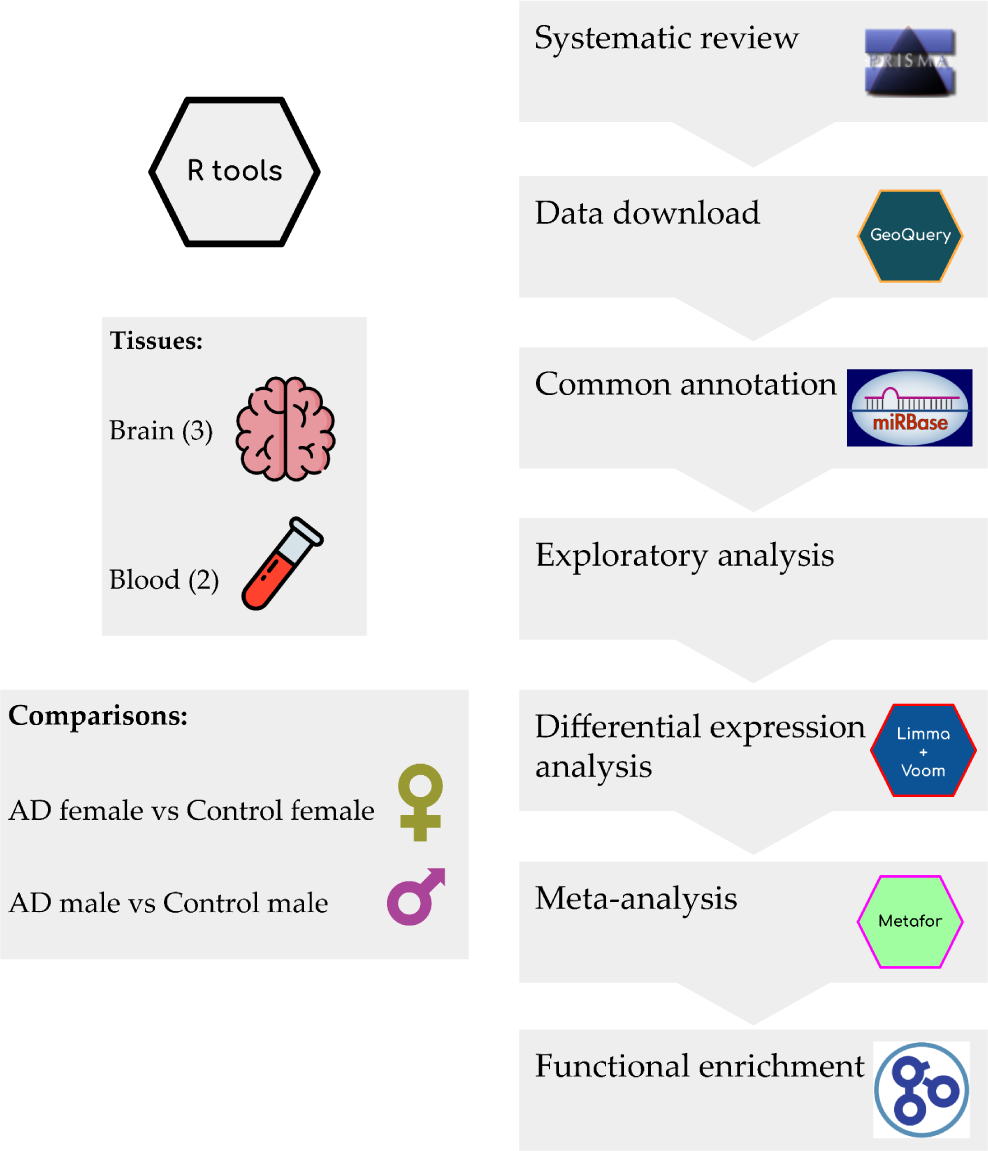
**Workflow and analysis design**. After data exploration and preprocessing, we retrieved relevant studies from the GEO-NCBI and ArrayExpress data repositories and performed differential miRNA expression analysis on each selected study. We performed four different meta-analyses (brain male, brain female, blood male, and blood female) and finally applied functional enrichment on the gene targets of the miRNAs identified in each meta-analysis

### Systematic Review

We developed a systematic review following PRISMA guidelines to identify suitable miRNA-focused expression studies in human AD. Studies must include information regarding the sex associated with each sample. Following the search criteria (**Figure 2**), we initially identified twenty-seven studies (twenty-six in GEO and one in ArrayExpress). Following the removal of duplicate studies (n=2), non-human studies (n=4), non-AD-focused studies (n=9), studies with non-suitable experimental designs (n=2), and non-miRNA-based studies (3), we selected seven studies as initially suitable for further analysis. Out of those seven studies, we left GSE63501 and GSE153284 out of the analysis due to the impossibility of access to the expression data and a lack of standardization, respectively. Therefore, we included five studies in the analysis: GSE157239 [47], GSE16759 [48], and GSE48552 [49] for brain tissue and GSE120584 [50] and GSE46579 [51] for blood samples **(Table 1)**.

**Figure 2.**
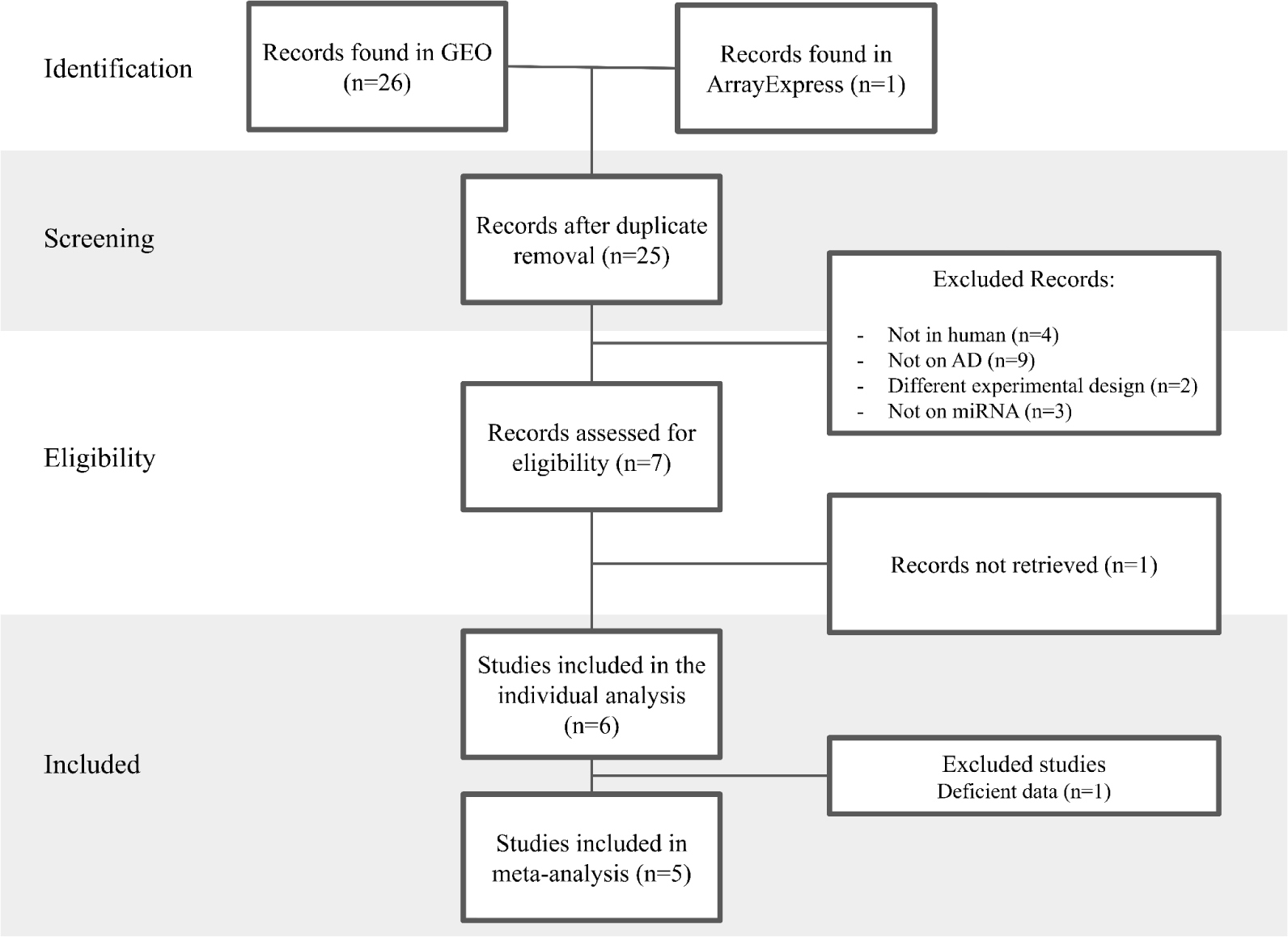
**Diagram of the systematic review based on PRISMA guidelines**

**Table 1.** Summary of the selected studies from the systematic review.

| ID | Study type | Platform | Samples | Control | Cases | Type of sample | PMID |
| --- | --- | --- | --- | --- | --- | --- | --- |
| **GSE157239** | Non-coding RNA profiling by array | GPL21572 | 16 | 8 | 8 | Brain (Temporal cortex) | 32920076 |
| **GSE16759** | Non-coding RNA profiling by array | GPL8757 | 8 | 4 | 4 | Brain (Parietal lobe) | 20126538 |
| **GSE48552** | Non-coding RNA profiling by high-throughput sequencing | GPL11154 | 12 | 6 | 6 | Brain (Prefrontal cortex) | 24014289 |
| **GSE120584** | Non-coding RNA profiling by array | GPL21263 | 1309 | 288 | 1021 | Blood (serum) | 34686734 |
| **GSE46579** | Non-coding RNA profiling by high-throughput sequencing | GPL11154 | 70 | 22 | 48 | Blood (whole blood) | 23895045 |

### Exploratory and Differential Expression Analysis

The exploratory analysis allowed an assessment of the distribution of the expression patterns in each study and to update the annotation of the miRNAs analyzed. We identified common miRNAs between studies and only considered those appearing in two or more studies for the integration analysis. Condition and sex distribution of the samples (**Figure 3**) skewed towards female patients in most studies, with a lower degree of male representation in control or AD groups. GSE120584 possessed a much higher sample size than the other selected studies (n=1309).

**Figure 3.**
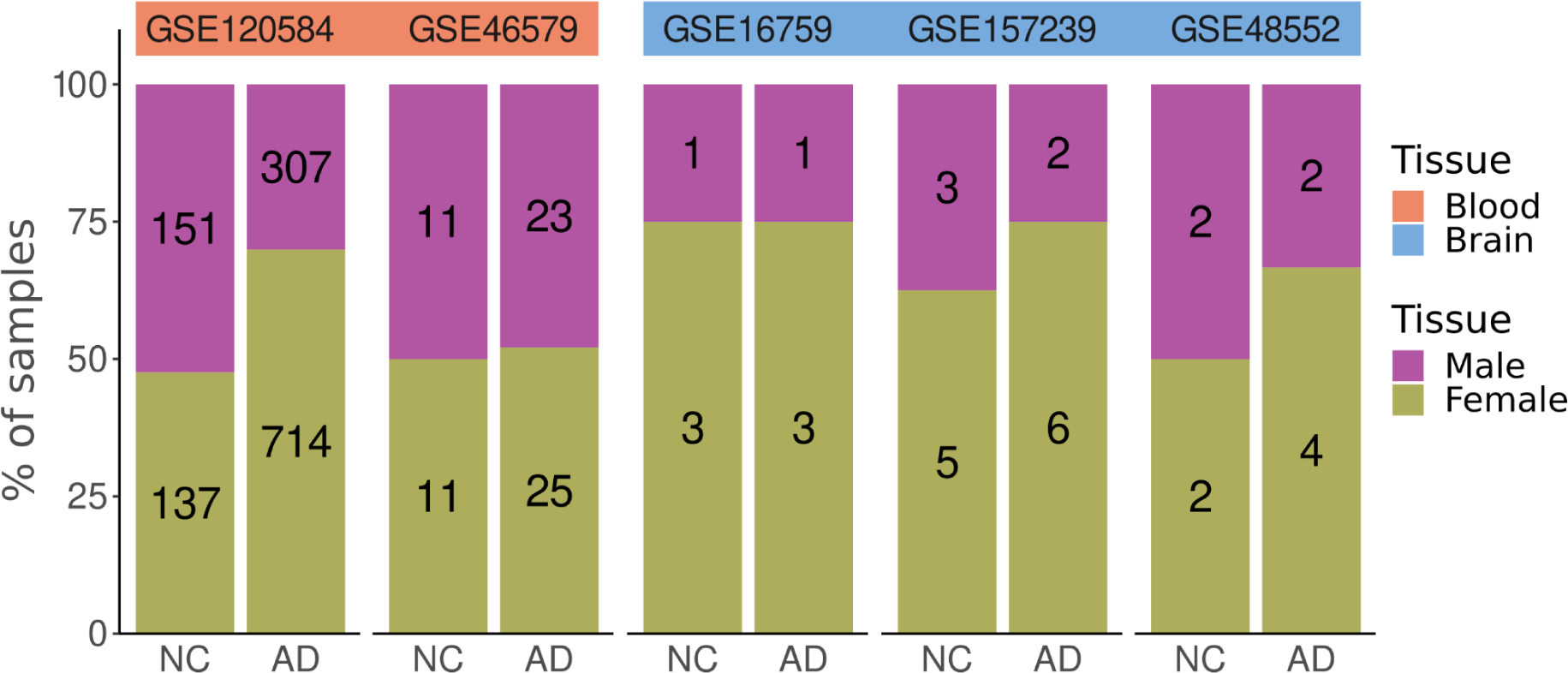
**Condition, tissue, and sex distribution of samples from selected studies**. NC, normal control; AD, Alzheimer’s disease

Expression data distribution did not provide evidence of anomalous samples, the hierarchical clustering of the samples did not display absolute divisions of the samples based on any of the experimental conditions, and the PCA visualization provided no evidence of bias.

The differential expression analysis of the individual studies returned profiles describing the altered expression of miRNAs in female and male patients in GSE120584, GSE46579, and GSE48552 studies but not in GSE16759 and GSE157239 **(Table 2)**. Subsequent analyses focused on integrating individual differential expression analysis to obtain robust results regarding the alteration of miRNAs for the comparisons considered.

**Table 2.**
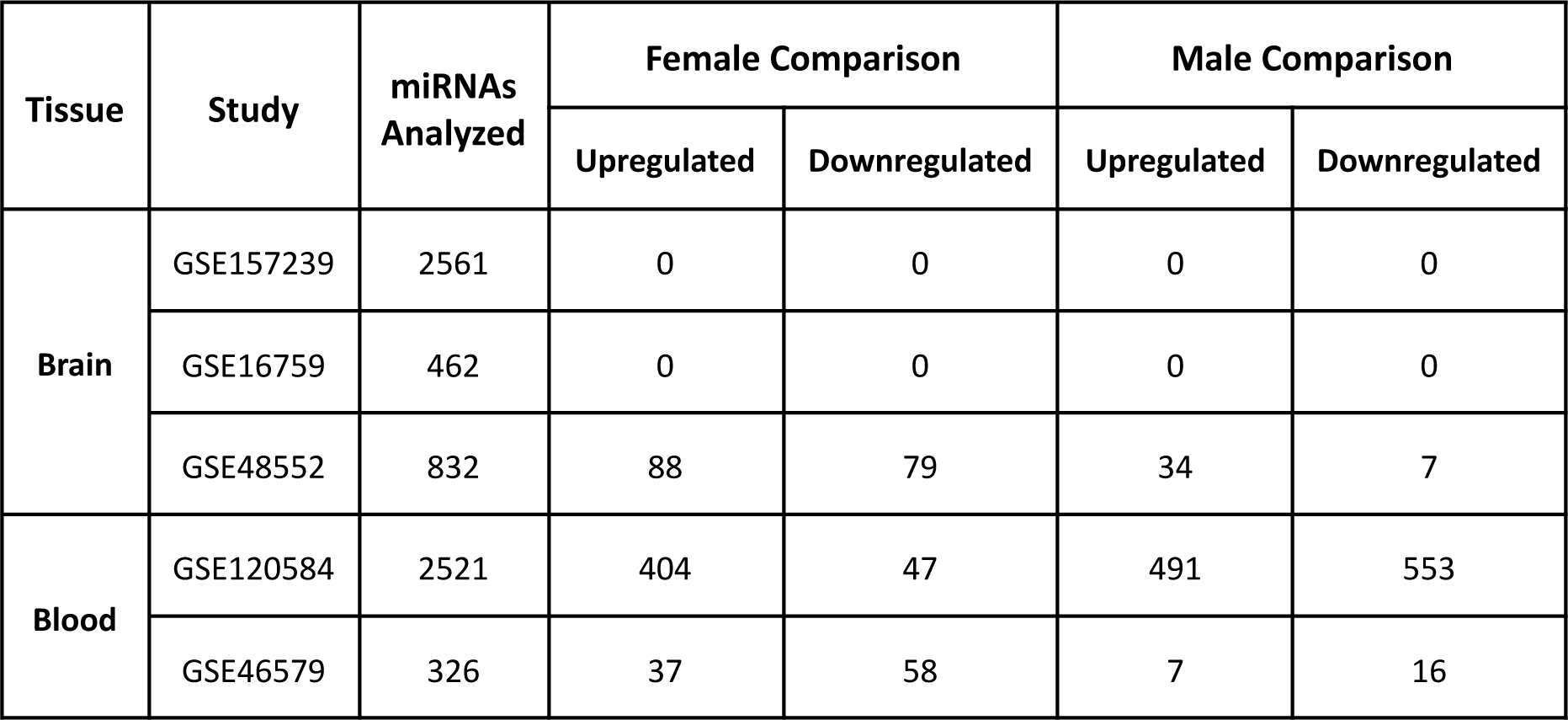
Differentially expressed miRNAs by study and by comparison.

| Tissue | Study | miRNAs Analyzed | Female Comparison |  | Male Comparison |  |
| --- | --- | --- | --- | --- | --- | --- |
|  |  |  | Upregulated | Downregulated | Upregulated | Downregulated |
| **Brain** | GSE157239 | 2561 | 0 | 0 | 0 | 0 |
|  | GSE16759 | 462 | 0 | 0 | 0 | 0 |
|  | GSE48552 | 832 | 88 | 79 | 34 | 7 |
| **Blood** | GSE120584 | 2521 | 404 | 47 | 491 | 553 |
|  | GSE46579 | 326 | 37 | 58 | 7 | 16 |

### Meta-analysis and Functional Enrichment

We performed four meta-analyses, integrating the differential miRNA expression results of sets of studies based on sample tissue (blood or brain) and sex (females and males) **(Table 3)**. We performed a random-effects DerSimonian-Laird (DL) meta-analysis on each combination’s differential miRNA expression results from the limma/limma+voom approach. We obtained a combined logFC for each miRNA under analysis and an associated BH-adjusted p-value.

**Table 3.**
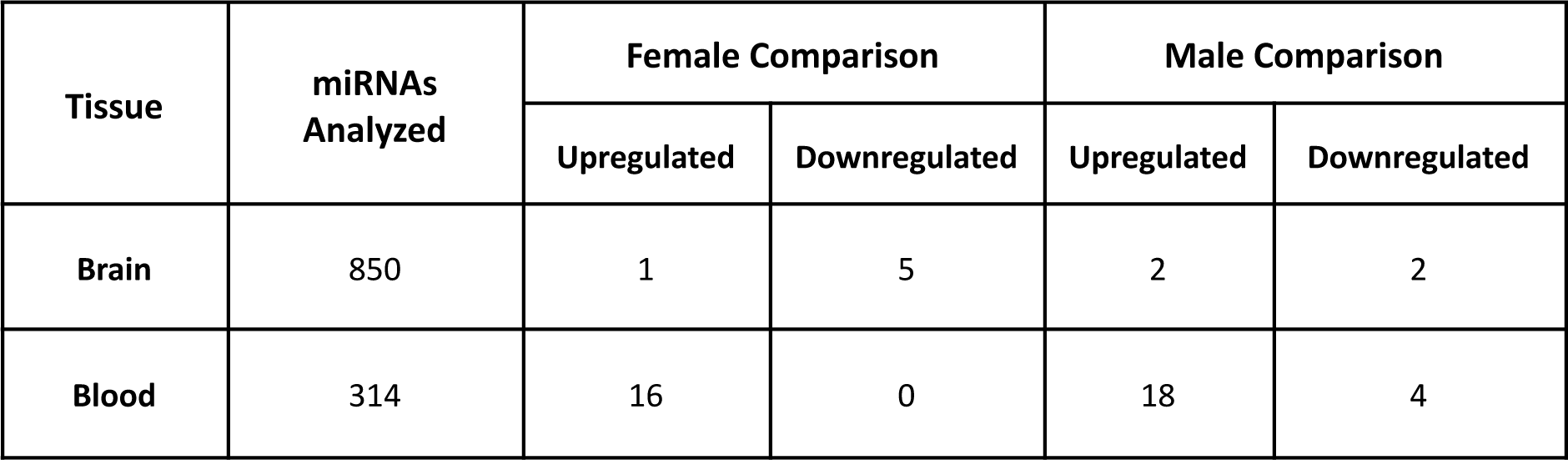
Results of differential expression meta-analyses based on individual studies.

| Tissue | miRNAs Analyzed | Female Comparison |  | Male Comparison |  |
| --- | --- | --- | --- | --- | --- |
|  |  | Upregulated | Downregulated | Upregulated | Downregulated |
| Brain | 850 | 1 | 5 | 2 | 2 |
| Blood | 314 | 16 | 0 | 18 | 4 |

### Blood meta-analyses

We found significantly altered miRNAs in female and male AD patients via meta-analyses based on blood samples (**Figure 4**). Sixteen miRNAs became significantly overexpressed in female AD patients compared to female control patients (**Figure 4A**), while eighteen miRNAs became significantly overexpressed and four underexpressed in male AD patients compared to male control patients (**Figure 4B**). We intersected profiles of overexpressed miRNAs in female and male AD patients for comparative purposes, which revealed a common increase in the expression of nine miRNAs in AD patients of both sexes and the exclusive overexpression of seven miRNAs in females and nine miRNAs in males (**Figure 4C**). Then, we compared the expression profiles of these altered miRNAs in male and female AD patients. miRNAs altered in female AD patients shared overall similar expression patterns in females and males (**Figure 4D**). miRNAs altered exclusively in male AD patients mainly shared similar expression patterns in males and females (**Figure 4E**); however, *hsa-mir-145-5p* displayed a significant decrease in males but did not change in females.

**Figure 4.**
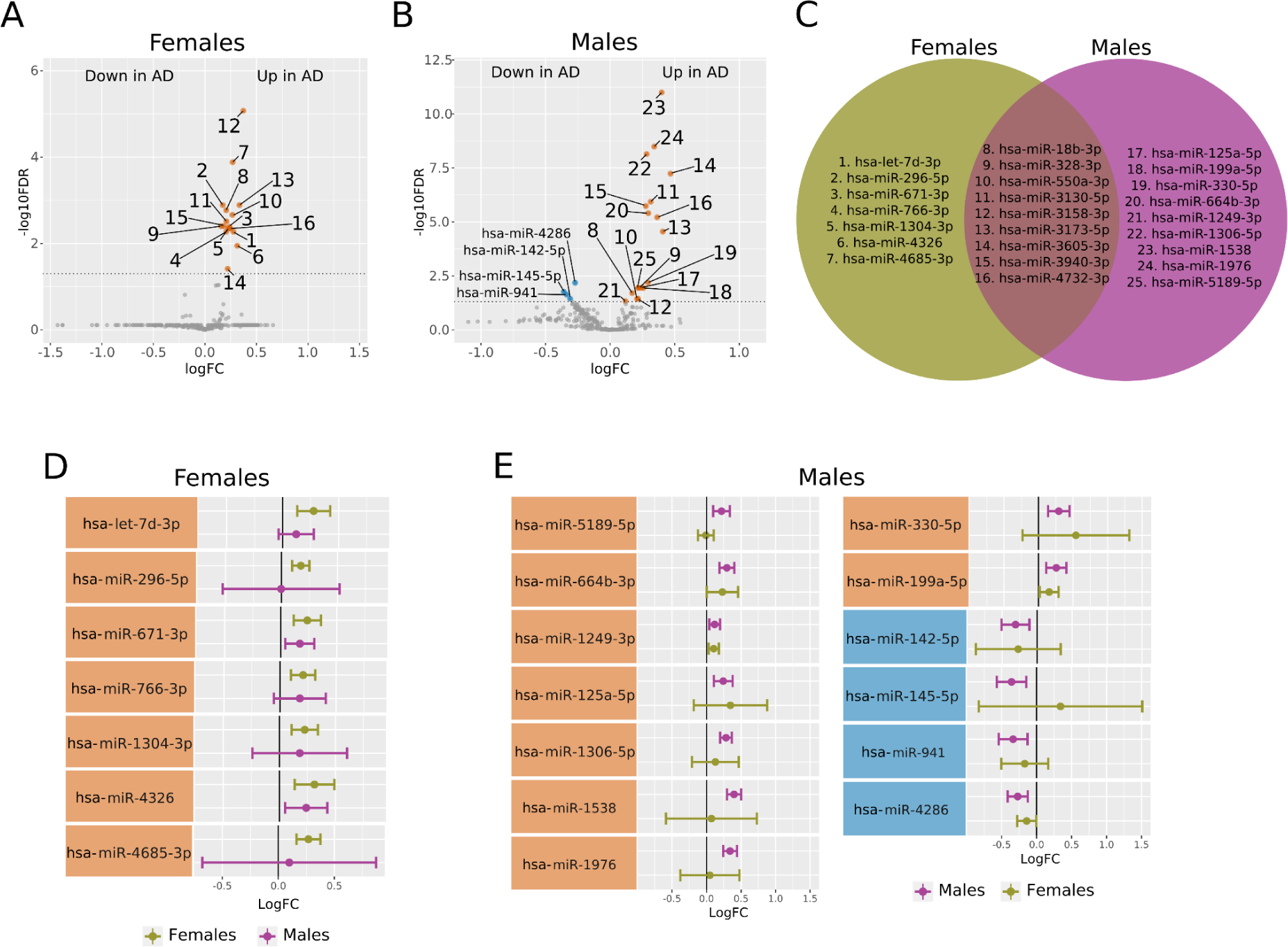
**Differential miRNA expression profiles in meta-analyses of blood samples from female and male AD and control patients**. **A.** Volcano plot showing overexpressed miRNAs (orange dots, sixteen miRNAs) in female AD patients. MiRBase IDs corresponding to displayed numbers are listed in **C**. Horizontal dashed gray line indicates -log_10_FDR (0.05). **B.** Volcano plot showing miRNAs underexpressed (blue dots, four miRNAs) and overexpressed (orange dots, eighteen miRNAs) in male AD patients. MiRBase IDs corresponding to displayed numbers are listed in **C**. Horizontal dashed gray line indicates -log10FDR (0.05). **C.** Venn diagram showing the intersection of overexpressed miRNAs in female and male AD patients. **D.** Plot comparing in both sexes the expression profiles of miRNAs exclusively altered in female AD patients. **E.** Plot comparing in both sexes the expression profiles of miRNAs exclusively altered in male AD patients

We next compared the target profiles of the differentially expressed miRNAs to unveil those genes that may be impacted by AD (**Supplementary Table 1**). The top genes targeted by miRNAs significantly altered in female AD patients included *ANKRD52* (target of eight miRNAs), *CELF1* and *LARP1* (target of seven miRNAs)*, CBX6, KMT2D, SETD5, SRCAP, SRRM2,* and *TAOK1* (target of six miRNAs). The top genes targeted by miRNAs significantly altered in male AD patients included *LARP1, FUS, BAZ2A, KMT2D,* and *DICER1*; the three miRNAs underexpressed in male AD patients targeted *BTBD3, NDN, NUP43, PIK3C2B, RAC1, RASA1, RCAN2, RNF38*, and *RPRM*.

Based on the complete miRNA transcriptomic profile, we performed a GSEA on the biological process (BP) ontology of GO terms (**Figure 5**) using gene targets of all miRNAs with altered expression in AD patients regardless of their directionality. This selection allows us to identify BPs affected by the upregulation and/or downregulation of distinct genes. The functional enrichment analysis in AD females revealed six altered BPs (**Figure 5A**); the ’protein polyubiquitination’ and ’response to transforming growth factor beta’ terms became downregulated while the ’positive regulation of cytoplasmic translation,’ ’cytoplasmic translation,’ ’mRNA splicing, via spliceosome’ and ’RNA splicing’ terms became upregulated. Meanwhile, we found 351 affected BP terms in male AD patients (**Supplementary Table 2**); nine BP terms mainly related to sensory perception and ion transmembrane transport increased, while 342 BP terms decreased. **Supplementary Table 3** and **Figure 5C** summarize the BPs affected and the top ten clusters and their parent terms, respectively.

**Figure 5.**
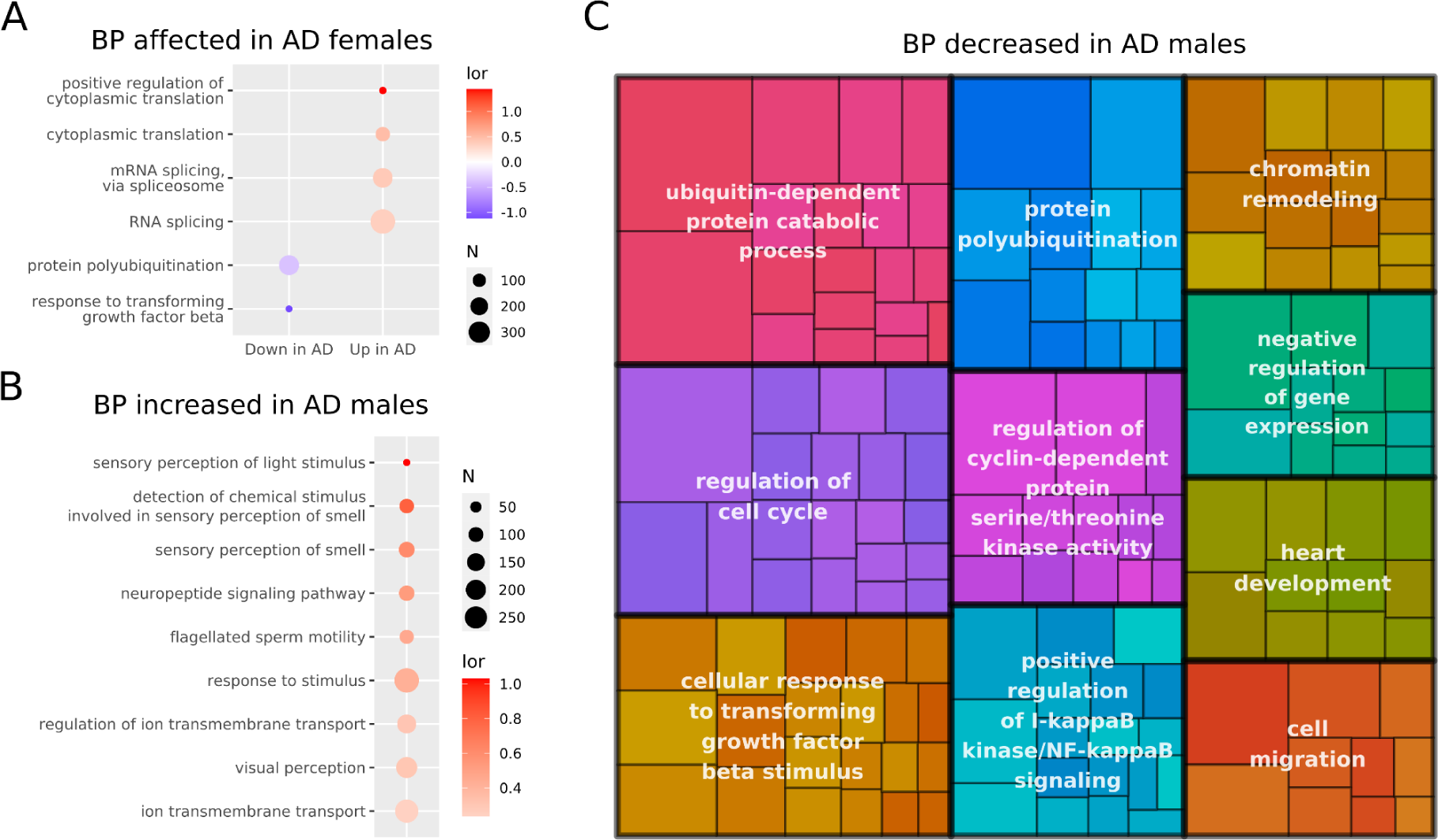
GSEA of blood meta-analyses in female and male AD and control patients. A. A dot plot describing the BP terms affected in female AD patients according to the gene targeted by miRNAs with significantly altered expression. Dots are colored based on log odds ratio (lor) value and their size are linked to the number of genes related to the BP. **B.** A dot plot describing the increased BP terms in male AD patients according to the gene targeted by miRNAs with significantly altered expression. Dots are colored based on log odds ratio (lor) value and their size are linked to the number of genes related to the BP. **C.** A tree map describing the top ten clusters of decreased BP terms in male AD patients according to the gene targeted by miRNAs with significantly altered expression

### Brain meta-analyses

The brain meta-analyses revealed the altered expression of miRNAs in female and male AD patients (**Figure 6**). Female AD patients displayed the underexpression of five miRNAs and the overexpression of one miRNA (**Figure 6A**); meanwhile, male AD patients displayed the underexpression of two miRNAs and the overexpression of two miRNAs (**Figure 6B**). The intersection of the altered miRNAs in female and male AD patients revealed five exclusively altered in females, three exclusively altered in males, and one common underexpressed miRNA (*hsa-miR-767-5p*). Those miRNAs altered exclusively in female AD patients shared similar expression patterns in both sexes except for *hsa-miR-105-3p*, which displayed an increase in female and a slight decrease in male AD patients (**Figure 6C**). Those miRNAs altered exclusively in male AD patients also shared similar expression patterns in females except for *hsa-mir-3149*, which displayed an increase in male and a non-significant decrease in female AD patients (**Figure 6D**).

**Figure 6.**
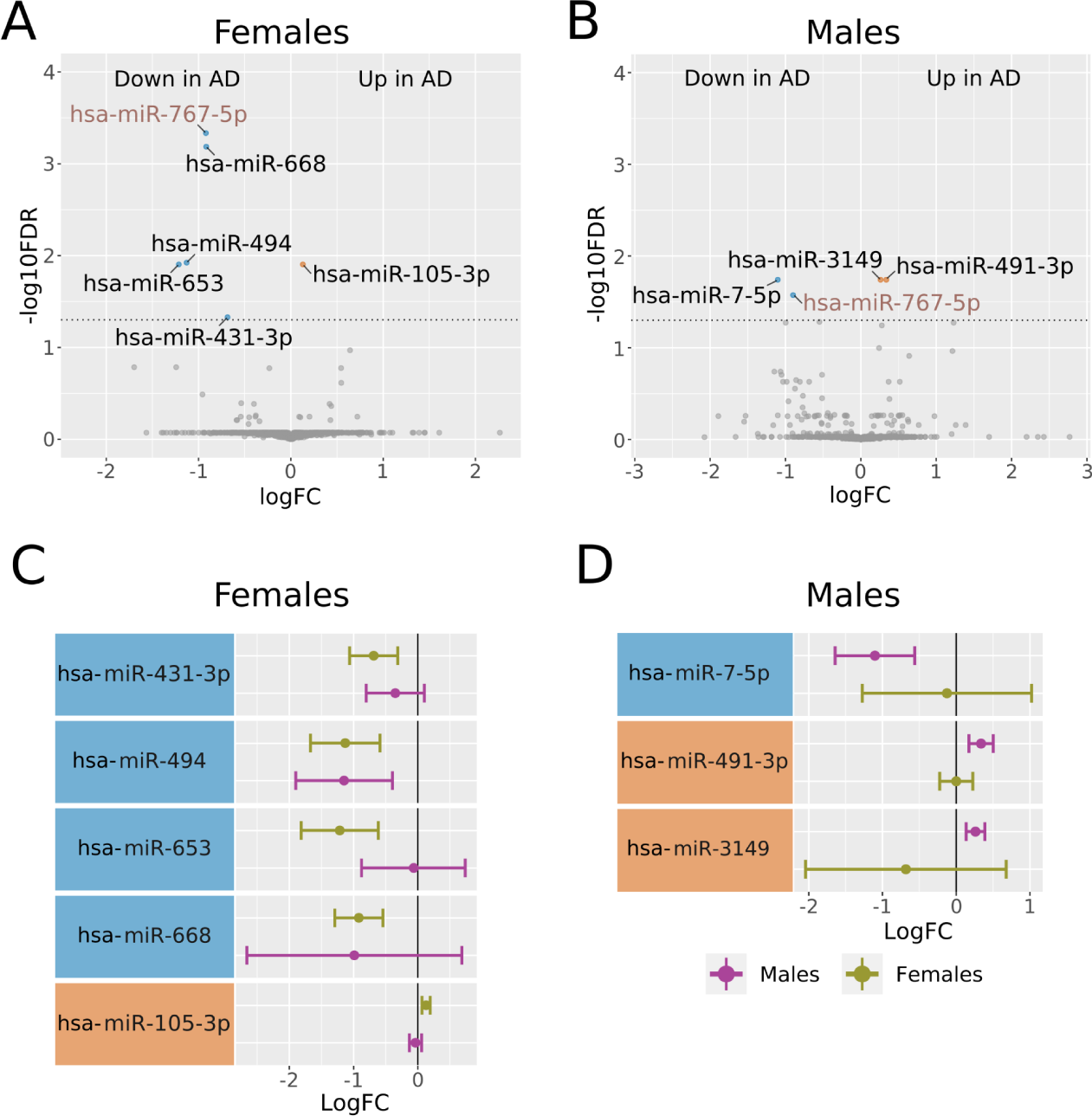
Differential miRNA expression profiles in meta-analyses of brain samples from female and male AD and control patients. A. Volcano plot showing miRNAs underexpressed (blue dots, five miRNAs) and overexpressed (orange dot, one miRNA) in female AD patients. The horizontal dashed gray line indicates -log_10_FDR (0.05). **B.** Volcano plot showing miRNAs underexpressed (blue dots, two miRNAs) and overexpressed (orange dot, two miRNAs) in male AD patients. The horizontal dashed gray line indicates -log_10_FDR (0.05). **C.** Plot comparing the expression profiles of miRNAs exclusively altered in female AD patients in both sexes. **D.** Plot comparing the expression profiles of miRNAs exclusively altered in male AD patients in both sexes

We then explored the target genes of the differentially expressed miRNAs in female and male AD patients (**Supplementary Table 4**). The genes targeted by the miRNA significantly increased in female AD patients (*hsa-miR-105-3p*) included *CBLN2, GOLIM4,* and *UHMK1* and the two miRNAs significantly decreased (*hsa-miR-431-3p* and *hsa-miR-767-5p*) included *MDM2, MTCH2,* and *MTRNR2L1*. The genes targeted by the two miRNAs significantly increased in male AD patients (*hsa-miR-491-3p* and *hsa-miR-3149*) included *NOM1 and ZNF226*, while the genes targeted by the two miRNAs significantly decreased represented a list of seventy-one genes. Notably, the functional enrichment performed on the miRNA profiles of female and male AD patients failed to reveal any specifically affected BP terms.

## Discussion

Despite the multiple sex-based differences described in AD symptomatology and epidemiology, their molecular basis remains unclear; furthermore, understanding the impact of sex-based differences on specific diseases remains crucial to improving clinical outcomes [52]. MiRNAs represent important regulators of gene expression whose relevance to AD has been underscored by recent studies explored [22,53,54]. We conducted a systematic review and four meta-analyses to unveil sex-based differences in miRNA profiles in the blood and brain of AD patients. The selected studies possessed a higher number of female AD samples, according to the epidemiology described [55]. Our results demonstrated similar alterations to miRNA expression profiles in the blood and brain of female and male AD patients. Moreover, miRNAs commonly affected in both sexes displayed a disease-associated increase and represent potential AD biomarkers. Finally, the functional enrichment analysis of miRNAs revealed sex-specific alterations of BP terms in the blood but not the brain.

### Blood meta-analyses

We found several deregulated miRNAs in female and male AD patients; the intersection analysis of these miRNAs points to the sex-specific alteration of miRNA expression, while miRNAs commonly affected in male and female AD patients represent potential disease biomarkers. We report a signature of seven miRNAs overexpressed in female AD patients compared to control females, with five previously unrelated to AD (*hsa-miR-296-5p, hsa-miR-766-3p, hsa-miR-1304-3p, hsa-miR-4326,* and *hsa-miR-4685-3p*). Previous studies reported the overexpression of *hsa-let-7d-3p* and *hsa-miR-671-3p* in AD patients in blood meta-analyses lacking a sex perspective [51,56]. Of the top targeted genes of significantly altered miRNAs in female AD patients, the nuclear speckle scaffold protein SRRM2 becomes accumulated in the cytoplasm of neurons in AD patients [57–59], while TAOK1 phosphorylation induces the formation of neurofibrillary tangles [60]. Notably, the top targeted genes primarily related to gene expression and chromatin organization rather than AD progression.

From the thirteen miRNAs specifically overexpressed by male AD patients, *hsa-miR-1306-5p* overexpression had been previously related to AD in a meta-analysis [61]. A study by Li et al. reported the downregulation of *hsa-miR-1306-5p* levels in the extracellular vesicles from serum samples of AD patients [62]; however, this finding did not preclude the detection of *hsa-miR-1306-5p* overexpression in blood as a component of circulating RNA. Additionally, Meng et al. reported that the expression of *hsa-miR-1306-5p* alleviated induced neurotoxicity in SK-N-SH cells treated with amyloid-β [63]. We also discovered a significant decrease in *hsa-miR-142-5p* expression in male AD patients. *Hsa-miR-142-5p* had been previously related to spatial learning and memory in AD animal models [64,65][65,66], suggesting a potential target for AD treatment.

We analyzed the functional effects of affected miRNAs based on their gene targets; a single miRNA can regulate the expression of multiple genes, resulting in complex interaction networks [67,68]. Female and male AD patients displayed divergent functional profiles affected by deregulated miRNA expression. In females, deregulated miRNAs in AD patients primarily altered splicing and translation. The dysregulation of tau splicing has been associated with neurodegenerative diseases and dementia [69], while altered translation could influence proteostasis and cytoplasmic protein accumulation, which significantly contributes to neuroinflammation and neurodegeneration [70,71]. In males, deregulated miRNAs in AD patients primarily increased BP terms related to smell/vision perception and ion transmembrane transportation. Notably, multiple regions involved in olfactory information processing display particular vulnerability in AD [72,73], suggesting odor identification as a potential but general biomarker of AD [74,75]. Meanwhile, Vitvitsky et al. reported the impairment of ion homeostasis in AD post-mortem brain samples [76]. We also observed decreased BP terms in male AD patients, including protein ubiquitination, which can influence the accumulation of misfolded proteins, hallmark of AD [77]. Furthermore, multiple BP clusters related to the regulation of gene expression through NF-κB signaling and chromatin remodeling [78] suggest a crucial role of gene expression misregulation in male AD patients.

### Brain meta-analyses

The brain meta-analyses revealed *hsa-miR-767-5p* as commonly overexpressed in female and male AD patients, in agreement with a previous report suggesting this miRNA as a biomarker candidate in the cerebrospinal fluid of AD patients [79]. No significantly deregulated miRNAs identified in female AD patients had previously reported links to AD except *hsa-miR-494,* which functions in stress pathways in AD [80]. The gene targets of affected miRNAs included the *MDM2* gene, which regulates p53 degradation and has previously reported links to AD [81]. The underexpression of *hsa-miR-7-5p* in male AD patients observed in this study is in contrast with the findings of a study by La Rosa et al., which reported increased *hsa-miR-7-5p* expression linked to the activity of the NLRP3 inflammasome [82]. *hsa-miR-491-3p* and *hsa-miR-3149*, both increased in the brains of male AD patients, lacked any previous link to AD; therefore, these miRNAs may represent sex-specific biomarkers.

### Strengths and limitations

We performed an in-silico strategy to evaluate and integrate the differential expression of miRNA transcriptomic studies. While previous systematic reviews and meta-analyses have been performed on this subject [61,83,84], to the best of our knowledge, we performed the first systematic review with sex as a central perspective to reveal links between AD and miRNA expression profiles in various tissues in female and male patients. Our approach allowed the analysis of differential miRNA expression profiles of males and females independently; however, we also highlighted a partial overlapping in miRNAs altered in both sexes (especially in blood samples).

Regarding potential limitations of the study, we found that the lack of information regarding the sex of individuals and the elevated number of studies conducted with only one sex restricted our sample size. Moreover, the disbalance between sexes in our sample size could represent a source of bias for the differential miRNA expression analysis. The selected studies also evaluated various brain regions, adding variability to the data due to highly heterogeneous cell populations in distinct areas. Finally, essential covariates such as medication usage, years of disease after diagnosis, and post-mortem interval were not included in the metadata of most original studies, thus increasing the levels of unexplained variability in the data.

### Perspectives and Significance

The results highlighted sex-based alteration to miRNA expression profiles in brain and blood samples from AD patients. We describe a panel of seven miRNAs that display altered expression in blood samples from female and male AD patients as potential disease biomarkers. We also observed sex-specific alterations in miRNA expression, highlighting the sex-based differential impact in AD of gene expression regulation and functional implications in multiple biological processes. Thus, the present study takes a novel approach to assess sex-based differences in miRNA expression in AD patients through a comprehensive bioinformatic strategy.

## Conclusions

In conclusion, our in-silico approach identified alterations in the expression of specific and common miRNAs in male and female AD patients that represent potential candidates as disease biomarkers, which is especially promising in blood samples as part of a liquid biopsy. Moreover, we identified sex-specific functional alterations associated with AD in blood samples related to RNA processing and translation in females and regulation of kinase activity, chromatin remodeling, and ubiquitination in males. These findings aim to foster a better understanding of miRNAs’ role in AD, emphasizing differences and similarities between males and females. Finally, we stress the critical role of open data sharing for scientific advancement.

## Declarations

### Ethics approval and consent to participate

Not applicable. **Consent for publication** Not applicable.

### Availability of data and materials

The data used for the analyses described in this work are publicly available at GEO [35]. The accession numbers of the GEO datasets downloaded are GSE16759, GSE46579, GSE48552, GSE120584, and GSE157239. The accession numbers of the ArrayExpress datasets downloaded are E-MTAB-1194 and E-MEXP-1416.

### Authors contributions

JLO and HC analyzed the data; FGG designed and supervised the bioinformatics analysis; HC and MRH designed and implemented the web tool; JLO, HC, and FGG wrote the manuscript; ZA, ISS, FG, MIV, FRG, AM, RM, BR and FGG helped in the interpretation of the results; HC, ZA, MIV, ISS, FRG, BR, AM, AP, MPM, RM and FGG writing-review and editing; FGG conceived the work. All authors read and approved the final manuscript.

## Acknowledgments

The authors thank the Principe Felipe Research Center (CIPF) for providing access to the cluster, co-funded by European Regional Development Funds (FEDER) in the Valencian Community 2014-2020. Irene Soler-Sáez thanks the Spanish Ministry of Universities for her predoctoral grant FPU20/03544. The authors also thank Stuart P. Atkinson for reviewing the manuscript.

## References

1. Reitz C, Rogaeva E, Beecham GW. Late-onset vs nonmendelian early-onset Alzheimer disease: A distinction without a difference? Neurol Genet. 2020;6:e512.

2. Scheltens P, De Strooper B, Kivipelto M, Holstege H, Chételat G, Teunissen CE, et al. Alzheimer’s disease. Lancet. 2021;397:1577–90.

3. Mendez MF. Early-Onset Alzheimer Disease. Neurol Clin. 2017;35:263–81.

4. Robinson M, Lee BY, Hane FT. Recent Progress in Alzheimer’s Disease Research, Part 2: Genetics and Epidemiology. J Alzheimers Dis. 2017;57:317–30.

5. Hauser S, Josephson S. Harrison’s Neurology in Clinical Medicine, 3E. McGraw Hill Professional; 2013.

6. Zhang X-X, Tian Y, Wang Z-T, Ma Y-H, Tan L, Yu J-T. The Epidemiology of Alzheimer’s Disease Modifiable Risk Factors and Prevention. J Prev Alzheimers Dis. 2021;8:313–21.

7. Kunkle BW, Grenier-Boley B, Sims R, Bis JC, Damotte V, Naj AC, et al. Genetic meta-analysis of diagnosed Alzheimer’s disease identifies new risk loci and implicates Aβ, tau, immunity and lipid processing. Nat Genet. 2019;51:414–30.

8. Soria Lopez JA, González HM, Léger GC. Alzheimer’s disease. Handb Clin Neurol. 2019;167:231–55.

9. Khan S, Barve KH, Kumar MS. Recent Advancements in Pathogenesis, Diagnostics and Treatment of Alzheimer’s Disease. Curr Neuropharmacol. 2020;18:1106–25.

10. Rajendran L, Paolicelli RC. Microglia-Mediated Synapse Loss in Alzheimer’s Disease. J Neurosci. 2018;38:2911–9.

11. Hampel H, Mesulam M-M, Cuello AC, Farlow MR, Giacobini E, Grossberg GT, et al. The cholinergic system in the pathophysiology and treatment of Alzheimer’s disease. Brain. 2018;141:1917–33.

12. Alzheimer’s Association. 2018 Alzheimer’s disease facts and figures. Alzheimers Dement. 2018;14:367–429.

13. Rocca WA. Time, Sex, Gender, History, and Dementia. Alzheimer Dis Assoc Disord. 2017;31:76–9.

14. Irvine K, Laws KR, Gale TM, Kondel TK. Greater cognitive deterioration in women than men with Alzheimer’s disease: a meta analysis. J Clin Exp Neuropsychol. 2012;34:989–98.

15. Hua X, Hibar DP, Lee S, Toga AW, Jack CR Jr, Weiner MW, et al. Sex and age differences in atrophic rates: an ADNI study with n=1368 MRI scans. Neurobiol Aging. 2010;31:1463–80.

16. Koran MEI, Wagener M, Hohman TJ, Alzheimer’s Neuroimaging Initiative. Sex differences in the association between AD biomarkers and cognitive decline. Brain Imaging Behav. 2017;11:205–13.

17. Buckley RF, Scott MR, Jacobs HIL, Schultz AP, Properzi MJ, Amariglio RE, et al. Sex Mediates Relationships Between Regional Tau Pathology and Cognitive Decline. Ann Neurol. 2020;88:921–32.

18. Dubois B, Feldman HH, Jacova C, Hampel H, Molinuevo JL, Blennow K, et al. Advancing research diagnostic criteria for Alzheimer’s disease: the IWG-2 criteria. Lancet Neurol. 2014;13:614–29.

19. Mattsson N, Lönneborg A, Boccardi M, Blennow K, Hansson O, Geneva Task Force for the Roadmap of Alzheimer’s Biomarkers. Clinical validity of cerebrospinal fluid Aβ42, tau, and phospho-tau as biomarkers for Alzheimer’s disease in the context of a structured 5-phase development framework. Neurobiol Aging. 2017;52:196–213.

20. Swarbrick S, Wragg N, Ghosh S, Stolzing A. Systematic Review of miRNA as Biomarkers in Alzheimer’s Disease. Mol Neurobiol. 2019;56:6156–67.

21. O’Brien J, Hayder H, Zayed Y, Peng C. Overview of MicroRNA Biogenesis, Mechanisms of Actions, and Circulation. Front Endocrinol . 2018;9:402.

22. Wei W, Wang Z-Y, Ma L-N, Zhang T-T, Cao Y, Li H. MicroRNAs in Alzheimer’s Disease: Function and Potential Applications as Diagnostic Biomarkers. Front Mol Neurosci. 2020;13:160.

23. Piscopo P, Bellenghi M, Manzini V, Crestini A, Pontecorvi G, Corbo M, et al. A Sex Perspective in Neurodegenerative Diseases: microRNAs as Possible Peripheral Biomarkers. Int J Mol Sci [Internet]. 2021;22. Available from: 10.3390/ijms22094423

24. Guo L, Zhong MB, Zhang L, Zhang B, Cai D. Sex Differences in Alzheimer’s Disease: Insights From the Multiomics Landscape. Biol Psychiatry. 2022;91:61–71.

25. Tarallo R, Giurato G, Bruno G, Ravo M, Rizzo F, Salvati A, et al. The nuclear receptor ERβ engages AGO2 in regulation of gene transcription, RNA splicing and RISC loading. Genome Biol. 2017;18:189.

26. Di Palo A, Siniscalchi C, Salerno M, Russo A, Gravholt CH, Potenza N. What microRNAs could tell us about the human X chromosome. Cell Mol Life Sci. 2020;77:4069–80.

27. Carè A, Bellenghi M, Matarrese P, Gabriele L, Salvioli S, Malorni W. Sex disparity in cancer: roles of microRNAs and related functional players. Cell Death Differ. 2018;25:477–85.

28. López-Cerdán A, Andreu Z, Hidalgo MR, Grillo-Risco R, Català-Senent JF, Soler-Sáez I, et al. Unveiling sex-based differences in Parkinson’s disease: a comprehensive meta-analysis of transcriptomic studies. Biol Sex Differ. 2022;13:68.

29. Català-Senent JF, Andreu Z, Hidalgo MR, Soler-Sáez I, Roig FJ, Yanguas-Casás N, et al. A deep transcriptome meta-analysis reveals sex differences in multiple sclerosis. Neurobiol Dis. 2023;181:106113.

30. Català-Senent JF, Hidalgo MR, Berenguer M, Parthasarathy G, Malhi H, Malmierca-Merlo P, et al. Hepatic steatosis and steatohepatitis: a functional meta-analysis of sex-based differences in transcriptomic studies. Biol Sex Differ. 2021;12:29.

31. Pérez-Díez I, Hidalgo MR, Malmierca-Merlo P, Andreu Z, Romera-Giner S, Farràs R, et al. Functional Signatures in Non-Small-Cell Lung Cancer: A Systematic Review and Meta-Analysis of Sex-Based Differences in Transcriptomic Studies. Cancers [Internet]. 2021;13. Available from: 10.3390/cancers13010143

32. Casanova Ferrer F, Pascual M, Hidalgo MR, Malmierca-Merlo P, Guerri C, García-García F. Unveiling Sex-Based Differences in the Effects of Alcohol Abuse: A Comprehensive Functional Meta-Analysis of Transcriptomic Studies. Genes [Internet]. 2020;11. Available from: 10.3390/genes11091106

33. Schulz J, Takousis P, Wohlers I, Itua IOG, Dobricic V, Rücker G, et al. Meta-analyses identify differentially expressed micrornas in Parkinson’s disease. Ann Neurol. 2019;85:835–51.

34. Page MJ, McKenzie JE, Bossuyt PM, Boutron I, Hoffmann TC, Mulrow CD, et al. The PRISMA 2020 statement: an updated guideline for reporting systematic reviews. BMJ. 2021;372:n71.

35. Barrett T, Wilhite SE, Ledoux P, Evangelista C, Kim IF, Tomashevsky M, et al. NCBI GEO: archive for functional genomics data sets--update. Nucleic Acids Res. 2013;41:D991–5.

36. Athar A, Füllgrabe A, George N, Iqbal H, Huerta L, Ali A, et al. ArrayExpress update - from bulk to single-cell expression data. Nucleic Acids Res. 2019;47:D711–5.

37. Rizzo ML. Statistical Computing with R, Second Edition. CRC Press; 2019.

38. Kozomara A, Birgaoanu M, Griffiths-Jones S. miRBase: from microRNA sequences to function. Nucleic Acids Res. 2019;47:D155–62.

39. Benjamini Y, Hochberg Y. Controlling the false discovery rate: A practical and powerful approach to multiple testing. J R Stat Soc. 1995;57:289–300.

40. DerSimonian R, Laird N. Meta-analysis in clinical trials. Control Clin Trials. 1986;7:177–88.

41. Viechtbauer W. Conducting Meta-Analyses inRwith themetaforPackage. J Stat Softw [Internet]. 2010;36. Available from: http://www.jstatsoft.org/v36/i03/

42. Gene Ontology Consortium. The Gene Ontology resource: enriching a GOld mine. Nucleic Acids Res. 2021;49:D325–34.

43. Ru Y, Kechris KJ, Tabakoff B, Hoffman P, Radcliffe RA, Bowler R, et al. The multiMiR R package and database: integration of microRNA-target interactions along with their disease and drug associations. Nucleic Acids Res. 2014;42:e133.

44. García FG. Métodos de análisis de enriquecimiento funcional de estudios genómicos. 2016.

45. Montaner D, Dopazo J. Multidimensional gene set analysis of genomic data. PLoS One. 2010;5:e10348.

46. Durinck S, Spellman PT, Birney E, Huber W. Mapping identifiers for the integration of genomic datasets with the R/Bioconductor package biomaRt. Nat Protoc. 2009;4:1184–91.

47. Henriques AD, Machado-Silva W, Leite REP, Suemoto CK, Leite KRM, Srougi M, et al. Genome-wide profiling and predicted significance of post-mortem brain microRNA in Alzheimer’s disease. Mech Ageing Dev. 2020;191:111352.

48. Nunez-Iglesias J, Liu C-C, Morgan TE, Finch CE, Zhou XJ. Joint genome-wide profiling of miRNA and mRNA expression in Alzheimer’s disease cortex reveals altered miRNA regulation. PLoS One. 2010;5:e8898.

49. Lau P, Bossers K, Janky R ’s, Salta E, Frigerio CS, Barbash S, et al. Alteration of the microRNA network during the progression of Alzheimer’s disease. EMBO Mol Med. 2013;5:1613–34.

50. Asanomi Y, Shigemizu D, Akiyama S, Sakurai T, Ozaki K, Ochiya T, et al. Dementia subtype prediction models constructed by penalized regression methods for multiclass classification using serum microRNA expression data. Sci Rep. 2021;11:20947.

51. Leidinger P, Backes C, Deutscher S, Schmitt K, Mueller SC, Frese K, et al. A blood based 12-miRNA signature of Alzheimer disease patients. Genome Biol. 2013;14:R78.

52. McGregor AJ, Hasnain M, Sandberg K, Morrison MF, Berlin M, Trott J. How to study the impact of sex and gender in medical research: a review of resources. Biol Sex Differ. 2016;7:46.

53. Lee CY, Ryu IS, Ryu J-H, Cho H-J. miRNAs as Therapeutic Tools in Alzheimer’s Disease. Int J Mol Sci [Internet]. 2021;22. Available from: 10.3390/ijms222313012

54. Arora T, Prashar V, Singh R, Barwal TS, Changotra H, Sharma A, et al. Dysregulated miRNAs in Progression and Pathogenesis of Alzheimer’s Disease. Mol Neurobiol. 2022;59:6107–24.

55. Aggarwal NT, Mielke MM. Sex Differences in Alzheimer’s Disease. Neurol Clin. 2023;41:343–58.

56. Takousis P, Sadlon A, Schulz J, Wohlers I, Dobricic V, Middleton L, et al. Differential expression of microRNAs in Alzheimer’s disease brain, blood, and cerebrospinal fluid. Alzheimers Dement. 2019;15:1468–77.

57. Lester E, Ooi FK, Bakkar N, Ayers J, Woerman AL, Wheeler J, et al. Tau aggregates are RNA-protein assemblies that mislocalize multiple nuclear speckle components. Neuron. 2021;109:1675–91.e9.

58. McMillan PJ, Strovas TJ, Baum M, Mitchell BK, Eck RJ, Hendricks N, et al. Pathological tau drives ectopic nuclear speckle scaffold protein SRRM2 accumulation in neuron cytoplasm in Alzheimer’s disease. Acta Neuropathol Commun. 2021;9:117.

59. Tanaka H, Kondo K, Chen X, Homma H, Tagawa K, Kerever A, et al. The intellectual disability gene PQBP1 rescues Alzheimer’s disease pathology. Mol Psychiatry. 2018;23:2090–110.

60. Giacomini C, Koo C-Y, Yankova N, Tavares IA, Wray S, Noble W, et al. A new TAO kinase inhibitor reduces tau phosphorylation at sites associated with neurodegeneration in human tauopathies. Acta Neuropathol Commun. 2018;6:37.

61. Yoon S, Kim SE, Ko Y, Jeong GH, Lee KH, Lee J, et al. Differential expression of MicroRNAs in Alzheimer’s disease: a systematic review and meta-analysis. Mol Psychiatry. 2022;27:2405–13.

62. Li F, Xie X-Y, Sui X-F, Wang P, Chen Z, Zhang J-B. Profile of Pathogenic Proteins and MicroRNAs in Plasma-derived Extracellular Vesicles in Alzheimer’s Disease: A Pilot Study. Neuroscience. 2020;432:240–6.

63. Meng S, Wang B, Li W. CircAXL Knockdown Alleviates Aβ-Induced Neurotoxicity in Alzheimer’s Disease via Repressing PDE4A by Releasing miR-1306-5p. Neurochem Res. 2022;47:1707–20.

64. Fu C-H, Han X-Y, Tong L, Nie P-Y, Hu Y-D, Ji L-L. miR-142 downregulation alleviates the impairment of spatial learning and memory, reduces the level of apoptosis, and upregulates the expression of pCaMKII and BAI3 in the hippocampus of APP/PS1 transgenic mice. Behav Brain Res. 2021;414:113485.

65. Liang W, Xie Z, Liao D, Li Y, Li Z, Zhao Y, et al. Inhibiting microRNA-142-5p improves learning and memory in Alzheimer’s disease rats via targeted regulation of the PTPN1-mediated Akt pathway. Brain Res Bull. 2023;192:107–14.

66. Zhang N, Gao Y, Yu S, Sun X, Shen K. Berberine attenuates Aβ42-induced neuronal damage through regulating circHDAC9/miR-142-5p axis in human neuronal cells. Life Sci. 2020;252:117637.

67. Ameres SL, Zamore PD. Diversifying microRNA sequence and function. Nat Rev Mol Cell Biol. 2013;14:475–88.

68. Barry G. Integrating the roles of long and small non-coding RNA in brain function and disease. Mol Psychiatry. 2014;19:410–6.

69. Andreadis A. Misregulation of tau alternative splicing in neurodegeneration and dementia. Prog Mol Subcell Biol. 2006;44:89–107.

70. Kurtishi A, Rosen B, Patil KS, Alves GW, Møller SG. Cellular Proteostasis in Neurodegeneration. Mol Neurobiol. 2019;56:3676–89.

71. Yu C-H, Davidson S, Harapas CR, Hilton JB, Mlodzianoski MJ, Laohamonthonkul P, et al. TDP-43 Triggers Mitochondrial DNA Release via mPTP to Activate cGAS/STING in ALS. Cell. 2020;183:636–49.e18.

72. Ohm TG, Braak H. Olfactory bulb changes in Alzheimer’s disease. Acta Neuropathol. 1987;73:365–9.

73. Braak H, Braak E. Frequency of stages of Alzheimer-related lesions in different age categories. Neurobiol Aging. 1997;18:351–7.

74. Wilson RS, Schneider JA, Arnold SE, Tang Y, Boyle PA, Bennett DA. Olfactory identification and incidence of mild cognitive impairment in older age. Arch Gen Psychiatry. 2007;64:802–8.

75. Tabert MH, Liu X, Doty RL, Serby M, Zamora D, Pelton GH, et al. A 10-item smell identification scale related to risk for Alzheimer’s disease. Ann Neurol. 2005;58:155–60.

76. Vitvitsky VM, Garg SK, Keep RF, Albin RL, Banerjee R. Na+ and K+ ion imbalances in Alzheimer’s disease. Biochim Biophys Acta. 2012;1822:1671–81.

77. Oddo S. The ubiquitin-proteasome system in Alzheimer’s disease. J Cell Mol Med. 2008;12:363–73.

78. Chen X-F, Zhang Y-W, Xu H, Bu G. Transcriptional regulation and its misregulation in Alzheimer’s disease. Mol Brain. 2013;6:44.

79. Dhawan A. Extracellular miRNA biomarkers in neurologic disease: is cerebrospinal fluid helpful? Biomark Med. 2021;15:1377–88.

80. Hojati Z, Omidi F, Dehbashi M, Mohammad Soltani B. The Highlighted Roles of Metabolic and Cellular Response to Stress Pathways Engaged in Circulating hsa-miR-494-3p and hsa-miR-661 in Alzheimer’s Disease. Iran Biomed J. 2021;25:62–7.

81. Proctor CJ, Gray DA. GSK3 and p53 - is there a link in Alzheimer’s disease? Mol Neurodegener. 2010;5:7.

82. La Rosa F, Mancuso R, Agostini S, Piancone F, Marventano I, Saresella M, et al. Pharmacological and Epigenetic Regulators of NLRP3 Inflammasome Activation in Alzheimer’s Disease. Pharmaceuticals [Internet]. 2021;14. Available from: 10.3390/ph14111187

83. Hu Y-B, Li C-B, Song N, Zou Y, Chen S-D, Ren R-J, et al. Diagnostic Value of microRNA for Alzheimer’s Disease: A Systematic Review and Meta-Analysis. Front Aging Neurosci. 2016;8:13.

84. Zotarelli-Filho IJ, Mogharbel BF, Irioda AC, Stricker PEF, de Oliveira NB, Saçaki CS, et al. State of the Art of microRNAs Signatures as Biomarkers and Therapeutic Targets in Parkinson’s and Alzheimer’s Diseases: A Systematic Review and Meta-Analysis. Biomedicines [Internet]. 2023;11. Available from: 10.3390/biomedicines11041113

